# Getting out of a mammalian egg: the egg tooth and caruncle of the echidna

**DOI:** 10.1101/2021.10.12.464116

**Authors:** Jane C Fenelon, Bennetts Abbie, Anthwal Neal, Michael Pyne, Stephen D Johnston, Alistair R Evans, Abigail S Tucker, Marilyn B Renfree

## Abstract

In the short-beaked echidna, *Tachyglossus aculeatus*, after an initial period of *in utero* development, the egg is laid in the pouch and incubated for 10 days. During this time, the fetuses develop an egg tooth and caruncle to help them hatch. However, there are only a few historical references that describe the development of the monotreme egg tooth. Using unprecedented access to echidna pre- and post-hatching tissues, the egg tooth and caruncle were assessed by micro-CT, histology and immunofluorescence, to map the changes at the morphological and molecular level. Unlike mammalian tooth germs that develop by invagination of a placode, the echidna egg tooth developed by evagination, similar to that of the first teeth in some reptiles. The egg tooth ankylosed to the premaxilla, rather than forming a mammalian thecodont attachment, with loss of the egg tooth post-hatching associated with high levels of odontoclasts, and apoptosis. The caruncle formed as a separate mineralisation from the adjacent nasal capsule, and as observed in birds and turtles, the nasal region epithelium expressed markers of cornification. Together, this highlights that the monotreme egg tooth shares many similarities with reptilian teeth, suggesting that this tooth is conserved from a common ancestor of mammals and reptiles.

## Introduction

The short-beaked echidna, *Tachyglossus aculeatus*, is one of five species in the mammalian subgroup Monotremata. The other living members of the order are the three long-beaked echidnas: *Zaglossus attenboroughi, Zaglossus bartoni* and *Zaglossus bruijni*, as well as the platypus, *Ornithorhynchus anatinus.* It is estimated that monotremes diverged from therian mammals (eutherians and marsupials) approximately 184 million years ago (Cúneo et al., 2013; Luo et al., 2011). Whilst the oldest known echidna fossils are 25 million years old, it is believed that echidnas diverged from platypuses between 55 and 70 million years ago (Davit-Béal et al., 2009; Zhou et al., 2021).

The fossil record shows that over time there has been a reduction in the tooth roots of platypus with the development of hardened keratinised palatal spines, which aid in mastication in modern monotremes (Davit-Béal et al., 2009; Musser & Archer, 1998). A number of platypus ancestor fossils have only a few teeth, showing the evolution away from tooth development and towards hardened pads (Archer et al., 1985; Pascual et al., 1992). In contrast, all echidna fossils identified to date are edentulous (Pascual et al., 1992). Since modern platypus nestlings still possess 3 cusped molars which are lost by adulthood, the echidna is assumed to have lost cheek teeth after diverging from the platypus. Monotremes have also lost some of the enamel forming genes over time and echidnas have lost additional enamel genes compared to platypuses, which still have a thin enamel layer on their cheek teeth (Luckett & Zeller, 1989; Zhou et al., 2021). This is consistent with other animals that evolved edentulism, such as pangolins, birds and baleen whales which have lost one or more enamel producing genes over time (Meredith et al., 2013; Meredith et al., 2014).

The only tooth which is retained in modern echidnas is the egg tooth, which is lost shortly after hatching. An ancestral amniote characteristic conserved in monotremes and reptiles that distinguishes them from therian mammals is oviparity. At the time the echidna egg is laid it is 15-17 mm in diameter and the embryo is at an early somite stage (Griffiths, 1968, 1989; Hughes & Hall, 1998; Semon, 1894). During the short incubation period, the embryo develops into a fetus and the young hatches from the egg after only 10-10.5 days. The monotreme eggshell is leathery, porous and consists of loosely wound keratinous fibres (Griffiths, 1968). Notably, monotremes develop both an egg tooth and a caruncle to escape from their egg (Griffiths, 1968; Hill & De Beer, 1950; Hughes & Hall, 1998).

Reptiles and birds also possess either a caruncle or an egg tooth. Reptiles, turtles, Rhynchocephalia and crocodilians have a caruncle, a thickened keratinised epithelium positioned above the nasal cartilages (Alibardi, 2020). In contrast, squamates have a true tooth that can be either single or paired (Fons et al., 20220). Even viviparous reptiles have an egg tooth, although it is smaller and hidden under a layer of connective tissue (Hermyt et al., 2017). The vast majority of birds also use an egg tooth to hatch out of their egg. However, the avian egg tooth is very similar to the caruncle of turtles and crocodiles, consisting of a sharp, keratinized ‘horn-like projection’ rather than being an actual tooth (Clark, 1961; Kingsbury et al., 1953; Wang et al., 2017). The structure and appearance of the avian egg tooth is incredibly uniform across most of the one hundred avian species examined so far (Wetherbee, 1959). Interestingly, the caruncle in monotremes is supported by a bony protrusion known as an os caruncle, and therefore, is very different from that observed in birds and reptiles (De Beer, 1949; Hill & De Beer, 1950). The relationship of the os caruncle to the premaxilla has been debated as to whether it is a just an extension of the premaxilla or is an independent ossification which fused with the premaxilla (Hill & De Beer, 1950).

Mammalian teeth have remained largely unchanged in structure throughout time (Davit-Béal et al., 2009). Tooth development in therian mammals is initiated by ligand-receptor interactions between the oral epithelium and mesenchyme during embryonic development (Tucker & Sharpe, 2004). The tooth itself forms in different stages called the bud, cap and bell stages (Tucker & Sharpe, 2004). First the oral epithelium thickens, and sinks into the neural crest derived mesenchyme, which condenses around the epithelium and creates the tooth germ; this process relies on both sonic hedgehog (SHH) and fibroblast growth factor (FGF) signals for proliferation and stratification of the dental placode (Li et al., 2016). The epithelium then extends into the mesenchyme, wrapping around the condensing mesenchyme to form the tooth cap (Tucker & Sharpe, 2004). The primary enamel knot forms at the tip of the tooth bud in the cap stage of development and acts as a signalling centre. The bell stage then follows this, where cyto-differentiation occurs and it is here that odontoblasts and ameloblasts form, which produce dentine and enamel respectively (Tucker & Sharpe, 2004).

Odontoblasts are columnar cells that form a uniform layer around the dental pulp cavity. They are differentiated from the dental pulp and odontoblastic secretions form the dentine layer on their outer surface (Balic & Thesleff, 2015). There are different types of dentine involved in tooth development including primary, secondary and tertiary dentine. Osteodentine is a type of dentine, named for its resemblance to bone, and is observed when tertiary dentine develops rapidly, and in doing so, traps odontoblasts and other nearby cells (Avery, 1994). After the development of the dentine layer, ameloblasts mature on the outer surface of the dentine and secrete proteins such as amelogenin leading to enamel formation between the ameloblasts and the dentine layer (Sire et al., 2007). Enamel is the hardest substance in the body and covers the tooth crown. After the development of this enamel layer the root of the tooth develops and the tooth erupts through the gums (Thesleff & Juuri, 2012).

The mechanism of attachment for most mammals is thecodonty, which refers to the presence of a tooth root being secured in a socket by a periodontal ligament (Spellerberg, 1982). Thecodonty is believed to be the ancestral tooth attachment mode in the most recent common amniote ancestor (Bertin et al., 2018). The mode of tooth attachment is a main point of difference between mammalian and reptilian teeth. The most common modes of tooth attachment in reptile species are acrodont and pleurodont, both of which involve fusion to the bone of attachment (Gans et al., 1985). The overall stages of tooth development and composition have remained largely unchanged throughout time between mammals and reptiles, with enamel knot like structures observed in some reptiles with complex teeth (Sulcova et al., 2020; Zahradnicek et al., 2014). In both mammals and reptiles, SHH is expressed in the early dental epithelium and acts to promote initiation of a tooth (Buchtova et al., 2008; Fons et al., 20220; Li et al., 2016).

The overall stages of egg tooth development in reptiles tends to also mirror that of other vertebrate teeth, depicted by a thickening of the oral epithelium and the bud, cap and bell stages, with development starting in the rostral part of the snout (Hermyt et al., 2017). The main difference is that the egg tooth develops earlier than all other dentition and, in the case of some snakes, is larger than any other tooth (Fons et al., 20220; Hermyt et al., 2017). There is, however, no mention of a root for the egg tooth, so the bone attachment may be the only support keeping the tooth in place.

Whilst there has been some research into the platypus egg tooth and caruncle (Green, 1930; Hill & De Beer, 1950), there are little data on how the echidna egg tooth and caruncle develop. The platypus egg tooth has been described as a pulp cavity surrounded by dentine and covered with an outer layer of presumptive enamel (Davit-Béal et al., 2009; Hill & De Beer, 1950), which is lost 2 days after hatching (Manger et al., 1998). In echidnas, the egg tooth is first observed as a median conical papilla protruding from the anterior end of the snout. Similar to the platypus, the echidna egg tooth is described as being attached to the premaxilla and containing a dental pulp covered in dentine (Hill & De Beer, 1950; Seydel, 1899). Here we have, therefore, focused on the echidna egg tooth and caruncle, to understand the potential homology of these structures with their functional equivalents across birds and reptiles, and to address issues such as method of attachment and mechanisms of loss.

## Results

### The fetal echidna egg tooth forms by evagination of the epithelium

To follow egg tooth development in the echidna post-oviposition fetal samples were imaged and sectioned. Externally, no egg tooth was visible in the day 4 fetus but by day 6 post-oviposition onwards, the egg tooth was clearly visible (Figure 1A-C). This was confirmed with the histology of the echidna fetal heads which showed substantial changes in growth, particularly in the tooth region, between day 4 and day 6 (Figure 1D, G). In the day 4 fetus, the epithelium was thickened and expressed SHH, confirming the egg tooth was at the placode stage of development (Figure 1F). No evidence of invagination of the placode into the underlying mesenchyme was evident but the mesenchyme under the placode had started to condense. At day 6 the egg tooth projected outwards forming a tooth by evagination, in contrast to the invagination observed during eutherian mammalian dental development (Figure 1G-H). SHH remained expressed in the evaginating epithelium, with an additional positive domain in the oral epithelium further back in the mouth (Figure 1I).

**Figure 1:**
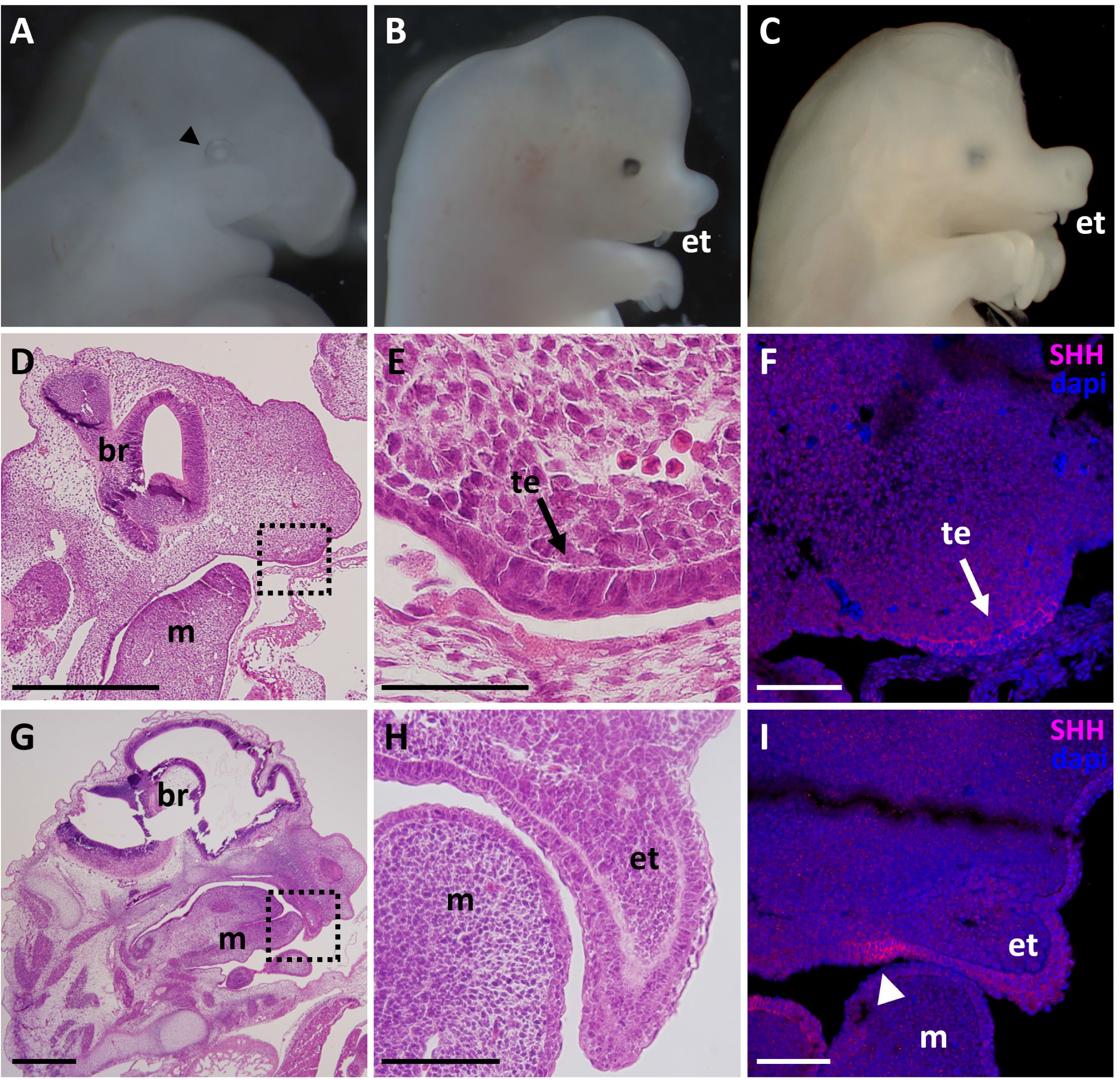
The echidna egg tooth forms by evagination of the epithelium. A-C: Photographs of post-oviposition echidna fetuses showing changes in external head and tooth morphology over time. A: day 4 fetus with no tooth or caruncle visible, black arrowhead indicates location of eye; B: by day 6.5 the egg tooth (et) is clearly visible; C: day 7 fetus; D: day 4 fetal head through the sagittal plane, the box indicates the magnified region in E; E: morphology of the thickened epithelium in the tooth region, F: SHH expression in the day 4 fetus showing significant expression in the oral thickened epithelium; G: day 6 fetal head through the sagittal plane, the box indicates the magnified region in H; H: morphology of day 6 fetal tooth, I: SHH expression in the day 6 fetus showing high expression in the oral epithelium (arrowhead). Blue staining= DAPI, red staining= SHH. br: brain, et: egg tooth, m: mandible, te: thickened epithelium. Scale bars F&I= 100 μm.

### The fetal echidna egg tooth forms a dentine layer continuous with the premaxilla bone

In the day 6 fetus, a mineralised layer had begun to form but the dental pulp had only just started to differentiate, and the cells were very densely packed (Figure 1H). By day 7.5, the dentine layer was clearly distinct with odontoblasts lining the inner surface, and the cells of the dental papilla appeared much more loosely packed (Figure 2A, E, I). The epithelium around the egg tooth was thickened at the tip with the formation of polarised cells that had a morphology similar to ameloblasts. However, an enamel layer was not evident at any of the stages examined (Figure 2K, L). The smooth layer of dentine at the tip of the egg tooth gave way to osteodentine at the base, with mesenchymal cells trapped within the dentine matrix (Figure 2I, J). This region appeared continuous with the premaxillary bone, as confirmed by Alizarin red staining (Figure 2D, H). No root was observed at any of the stages examined. The egg tooth is therefore, anchored directly to the bone.

**Figure 2:**
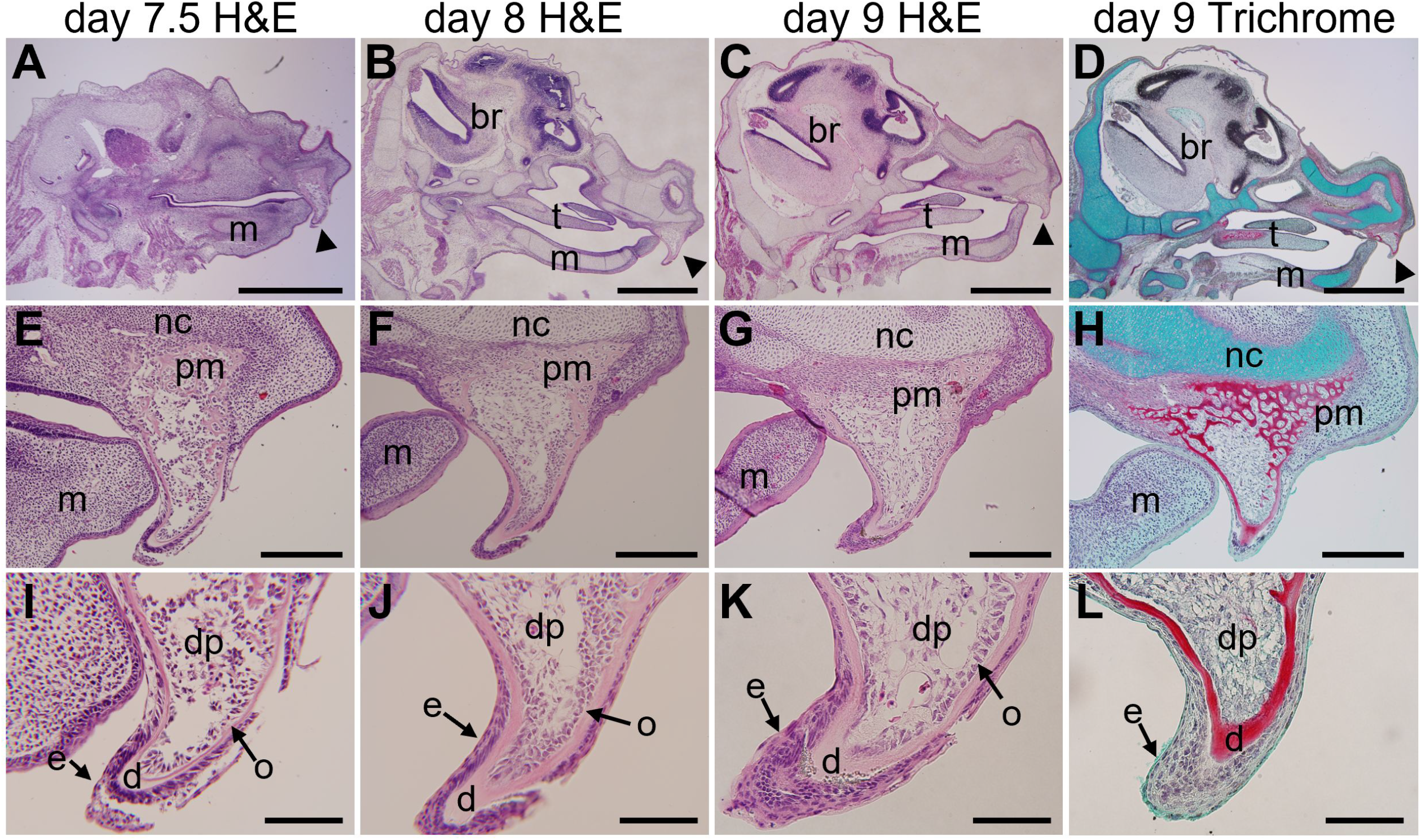
The echidna egg tooth forms a dentine layer continuous with the premaxilla bone. A: day 7.5 fetal head through the sagittal plane, B: day 8 fetal head through the sagittal plane, C: day 9 fetal head through the sagittal plane, D: trichome stained day 9 fetal head through the sagittal plane, E-H: higher power image of A-D showing the continuous dentine layer between the tooth and premaxilla, I-L: high power magnification of the tooth region in E-H. br: brain, dp: dental papilla, d: dentine, e: epithelium, et: egg tooth, m: mandible, t: tongue, o: osteoblasts. Arrows indicate the developing tooth region. All images H&E stained unless otherwise indicated. Trichrome stained sections: bone and dentine = red; cartilage =blue. Scale bars A-D: = 1 mm, E-H: 200 μm, I-K: 100 μm.

### The premaxilla unites the egg tooth and caruncle

The caruncle was first detected in the day 7.5 fetus and was clear in the day 8 fetus as an ossified circle of mesenchyme pressed against the nasal capsule (Figure 3A, B). Unlike the nasal capsule, which expresses high levels of Type II collagen, the caruncle did not express this cartilage marker, and stained with alizarin red, suggesting it is indeed a bone (Figure 3B, D, E, I). The caruncle could be seen to attach directly with the premaxillary bone, which extended upwards over the nasal cartilages, thereby, uniting the egg tooth and caruncle (Figure 3G, H). To confirm the arrangement of these skeletal elements, micro-CT was used to show the tissue in 3D (Figure 3C, F Supplementary Figure 3). The dentine of the egg tooth could be observed at day 8 connecting to the trabecular bone of the premaxilla, with two arms of bone extending up from the premaxilla to contact the caruncle (Figure 3F). The development of the premaxilla was more advanced than the other bones of the head, highlighting the need for this region to develop in advance of the rest of the structures to support the egg tooth and caruncle.

**Figure 3:**
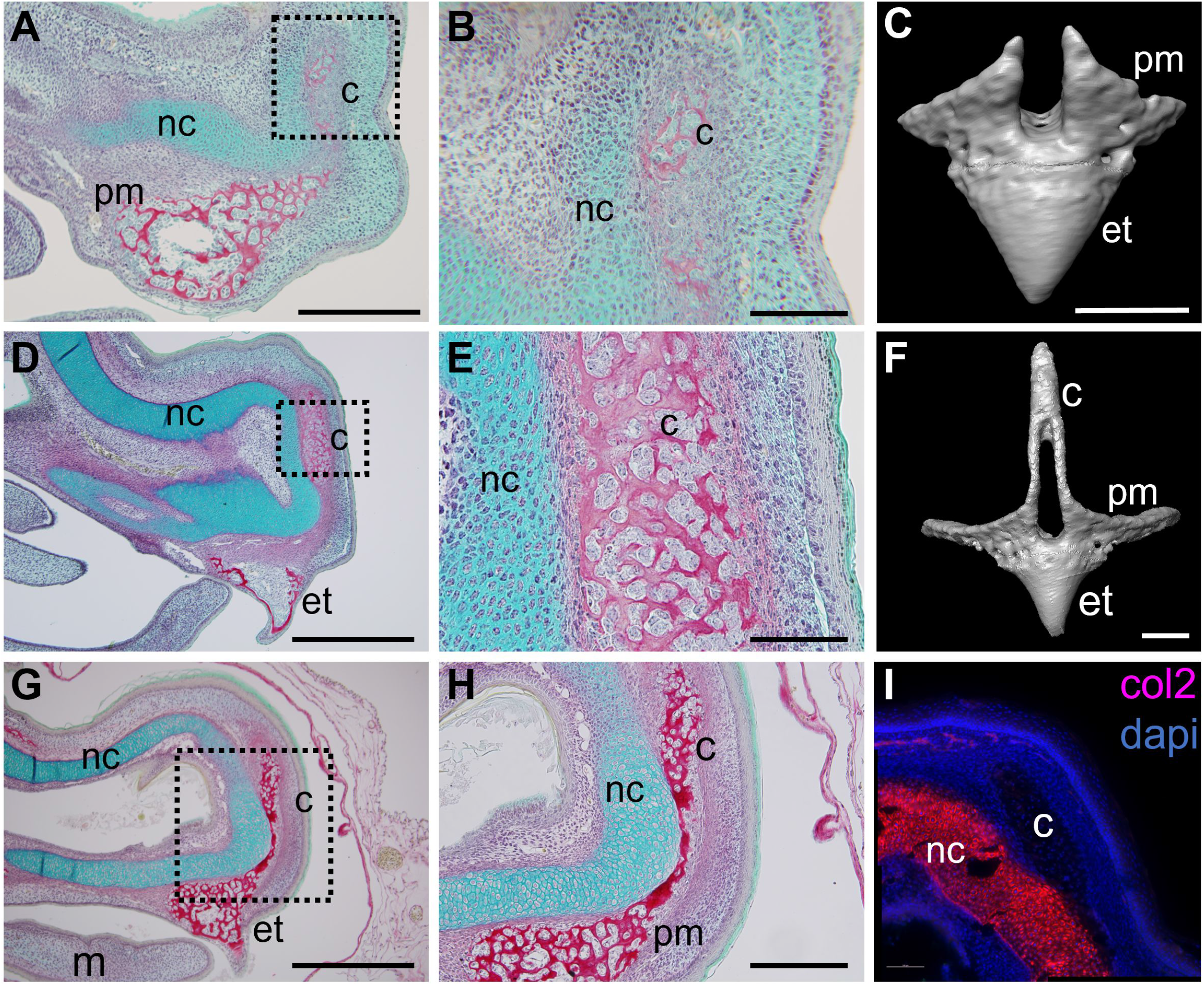
The premaxilla unites the egg tooth and the caruncle. A: Trichrome stained egg tooth and caruncle of a day 7.5 fetus; B: The initial calcification of the caruncle; C: Micro-CT volume rendering of the premaxilla and egg-tooth region in the same fetus; D: Trichrome stained egg tooth and caruncle of a day 9 fetus; E: The trabecular bone of the caruncle; F: Frontal plane view of a micro-CT volume rendering of the mineralised structures of the egg tooth, caruncle and premaxilla in a day 8 fetus; G: The snout of a mature fetus showing the egg tooth sharing the same attachment to the premaxilla as the caruncle; H: The connection between the caruncle and the egg tooth via the trabecular bone of the premaxilla; I: Immunofluorescence of collagen type II in the day 2 pouch young. Dotted boxes indicate area of magnified regions in B, E & H. c: caruncle, et: egg tooth, nc: nasal cartilage, pm: premaxilla. Trichrome stained sections: bone and dentine = red; cartilage =blue. Scale bars: A, F, H 200 μm, B & E: 100 μm, C: 50 μm, D & G: 500 μm. Figure supplement 3 shows the microCT scans of the additional egg tooth structures.

The epithelium overlying the osseous caruncle was thickened compared to the epithelium over the rest of the head as has been previously reported (Figure 4A) (Hill & De Beer, 1950). To observe the pattern of differentiation in more detail, markers of the different layers of epithelium were investigated. Keratin 14 is often used as a marker of the basal epithelium and is expressed in mammalian skin from early stages of development as the cells commit to stratification (Koster & Roop, 2004). Around the head, Keratin 14 was expressed in the basal layer of the epithelium, however, in the caruncle it was expressed in the intermediate layer (Figure 4B, C). Keratin 8, another marker of uncommitted epithelium (Liu et al., 2013), was also expressed in the head epithelium but was absent from the caruncle (Figure 4D, E). Finally, Loricrin, a marker of terminal differentiation, was only observed in the caruncle epithelium, in the top layer (Figure 4B), confirming the caruncle epithelial cells had formed a cornified layer.

**Figure 4:**
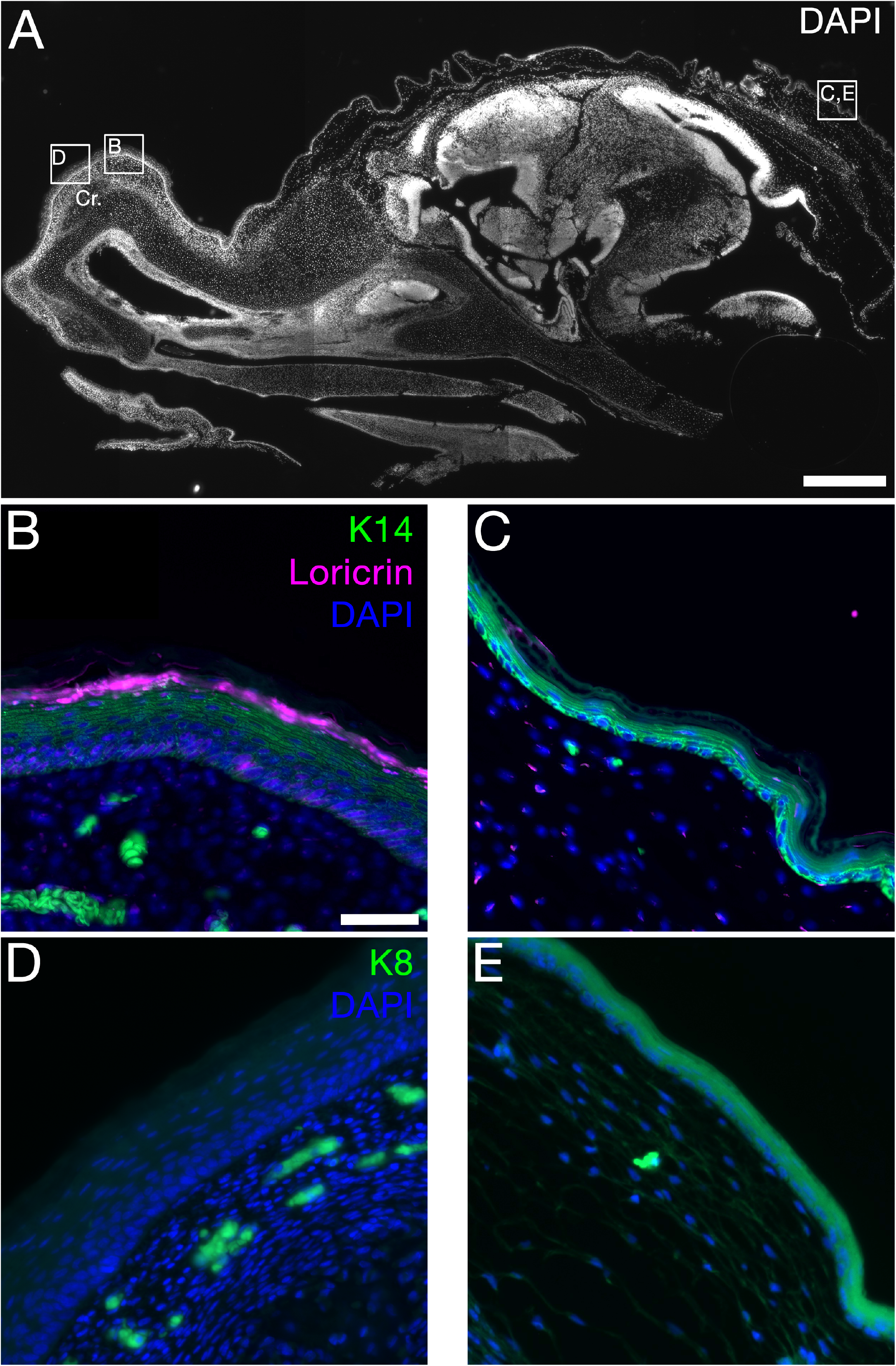
Precocious stratification of the ectoderm over the caruncle. A: DAPI stained Sagittal section through echidna head at day 0 pouch young. Boxes indicate locations shown in B-E. B,C: Expression by immunofluorescence in green of cytokeratin 14 (K14) and in magenta of Loricrin in the epidermal epithelium of the caruncle (B) and the back of the cranium (C). The epithelium at the caruncle is highly stratified compares to that at the back of head. K14 is expressed in the intermediate layer of the stratified epithelium at the caruncle (B), and strongly expressed in the basal layer at the back of the head (C). Loricrin, a marker of terminally differentiated epidermis, is expresses at the caruncle (B), but is absent form the back of the head (C). D,E: Immunofluorescence in green expression of cytokeratin 8 (K8), a marker of immature epidermal epithlium. K8 is not expressed in at the caruncle (D) but is expressed in the epithelium surrounding the back of the head (E). C: caruncle. Scale bars: A= 500 μm, B-E= 50 μm

### Post-hatching loss of the egg tooth by apoptosis and odontoclast activity

Loss of the egg tooth was followed by micro-CT post-hatching (Figure 5A-C). The egg tooth, clearly evident on the day of hatching was lost in the day 4 pouch young but was evident in some specimens at day 3 (Figure 5B-C). The mechanism of loss was therefore followed by TRAP staining to identify clast cells and by TUNEL for apoptosis. At the day of hatching very few TRAP positive cells were evident in the egg tooth and surrounding tissue (Figure 5D). In contrast at day 2, just before loss of the tooth, high numbers of TRAP positive cells were evident, particularly within the tooth against the dentine layer (Figure 5E). Thinning and breaks in the dentine were evident, suggesting that the tooth was removed by odontoclasts. High levels of apoptotic cells, as labelled with TUNEL, were evident in the dental pulp, particularly at the border with the premaxilla, highlighting loss of this tissue after hatching (Figure 5F). A combination of detachment of the dentine and death of the cells of the pulp therefore led to its loss. This mode of loss was also observed in the platypus, where breaks in the osteodentine were evident, flanked by multinucleated clast cells (Supplementary Figure 5). Multinucleate clasts cells were also identified in the dental papilla up against the dentine, suggesting that this mode of removal from the inside is conserved in monotremes (Supplementary Figure 5C). In contrast to the egg tooth, no TRAP staining was evident in the echidna caruncle up until day 4 pouch young (data not shown). In keeping with this, micro-CT analysis indicated that the caruncle was still present in a day 11 pouch young but was no longer evident by approximately day 50 (Figure 5 G-I).

**Figure 5:**
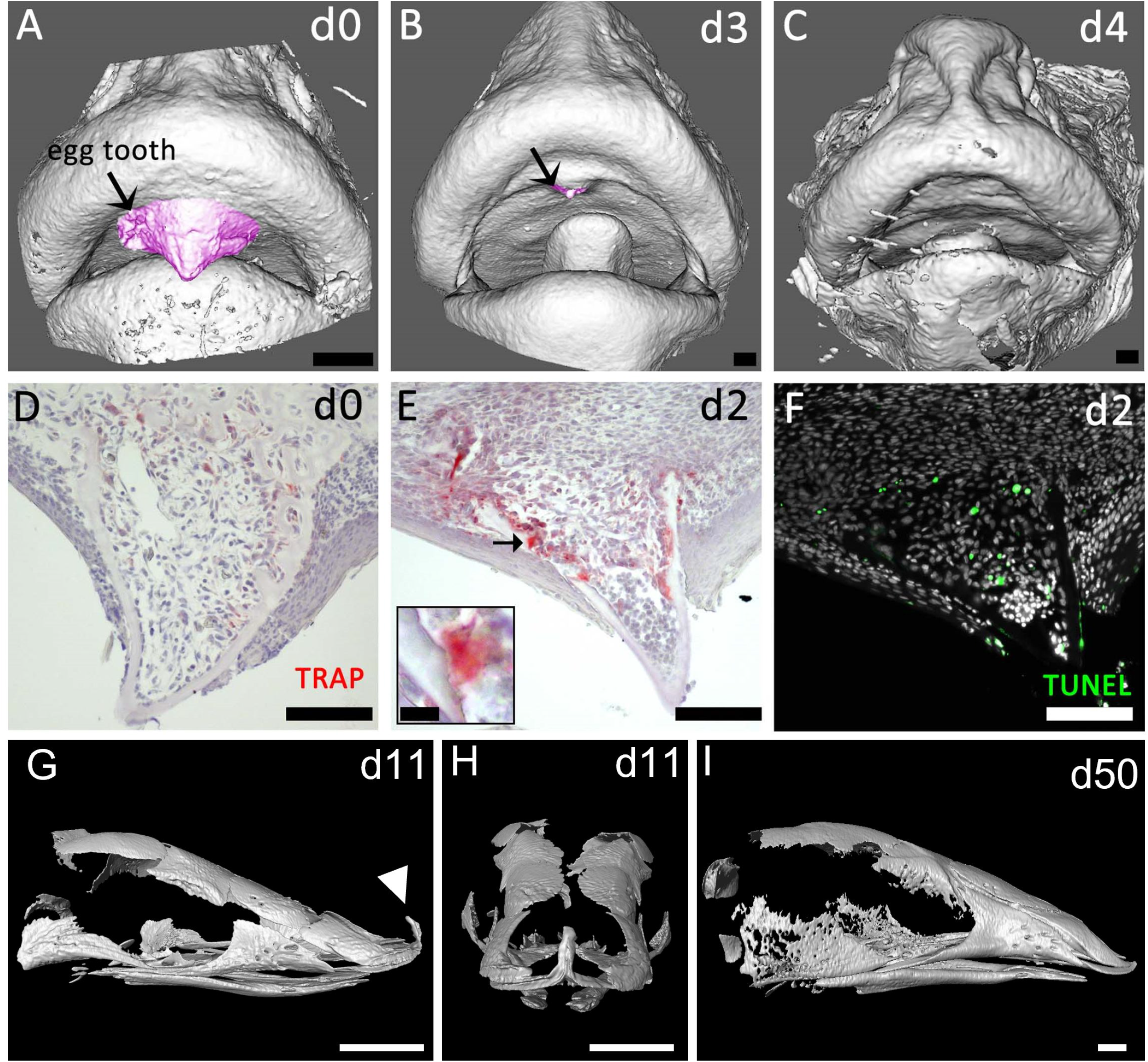
Post-hatching loss of the egg tooth by apoptosis and odontoclast activity. A-C: Soft tissue contrasted micro-CT volume rendering of the mouth region in d0 (A), d3 (B) and d4 (C) pouch young, showing the egg tooth (highlighted in pink) present at d0 and d3, but not at d4. D,E: TRAP staining showing no odontoclast activity in the egg tooth of d0 pouch young (D), whereas at d2 (E) odontoclasts are acting on the inner surface of the egg tooth dentin. Arrow in E shows location of high magnification insert demonstrating odontoclast activity associated with a break in the egg tooth dentin. F: Fluorescence TUNEL staining shows high levels of apoptosis in the pulp of the egg tooth of d2 pouch young. G-I: Mineralised tissue micro-CT volume rendering of the skull in d11 (G-H) and approximately d50 (I) pouch young showing continuation of the caruncle (white arrow), whereas it is gone by d50. Scale bars: A-C = 500 μm, D-F = 100 μm, inset E = 10 μm, G-I = 2mm. Figure Supplement 5 shows the beginnings of similar detachment in the platypus, *Ornithorhynchus anatinus* at 2 days post-hatching.

## Discussion

Echidna fetuses were found to undergo the most significant development with regards to tooth development between day 4 and day 7.5 post-oviposition, with SHH expressed in the thickening oral epithelium at day 4 and with the tooth reaching maturity by day 7.5. The finding of SHH in the egg tooth placode is consistent with the conserved expression of SHH in both mammals and reptiles in defining the position of the dental lamina and promoting tooth development (Balic & Thesleff, 2015; Buchtova et al., 2008). SHH expression was particularly high in the day 6 fetus where the tooth meets the mouth, which may indicate the location of a signalling centre controlling the projection of the tooth or the presence of an additional dental placode that arrests development at this stage.

Interestingly, instead of the placode invaginating into the surrounding mesenchyme, as observed in eutherian mammals, the echidna placode protruded out of the mouth. This evagination is also observed in lizards and crocodilians as part of the development of the first set of teeth (Westergaard & Ferguson, 1990; Zahradnicek et al., 2012). Evaginating tooth-like structures have also been observed in avian talpid mutants (Harris et al., 2006). While the later developing teeth in reptiles form from a dental lamina deep within the jaw, the first null generation teeth form superficially and project out of the mouth (Westergaard & Ferguson, 1990; Zahradnicek et al., 2012). These early teeth are lost by resorption or are expelled from the mouth and are often made of dentine without an enamel coating (Zahradnicek et al., 2012). Interestingly, evaginating teeth develop in some mouse mutants after manipulation of Wnt signalling, highlighting that a simple switch may be able to convert evaginating to invaginating tooth germs (Kim et al., 2021). The similarities in development of the echidna egg tooth and the evaginating early teeth in reptiles and archosaurs, could suggest that they are homologous, with this method of tooth development conserved from the last common amniote ancestor of synapsids and sauropsids.

The mature echidna egg tooth at hatching was comprised of layers of dental pulp, odontoblasts and dentine. However, whilst the odontoblasts were comprised of a clearly defined layer, they were more irregular in shape compared to the uniformly columnar odontoblasts typically identified in mammalian teeth. These more rudimentary odontoblasts are likely due to the temporary nature of the tooth. The dentine layer developed rapidly. Whilst the egg tooth was not fully mineralised at day 6, it was at its maximum thickness and mineralised only 1.5. days later. Histological examination of the dentine also revealed that it contained trapped cells at the base of the tooth, which resembled trabecular bone making it difficult to distinguish where the tooth ended and the premaxilla started. We identified this layer as a type of dentine known as osteodentine which can superficially resemble bone due to its rapid formation that can cause cells to be trapped inside it (Avery, 1994). This is consistent with the rapid formation of the echidna egg tooth and the rudimentary development of the odontoblasts.

While most of the cell types visible in mammalian teeth were observed in the mature echidna egg tooth, no enamel could be identified in histological sections of the tooth at any stage. Previous platypus egg tooth studies had suggested that there may be a layer of presumptive enamel around the egg tooth and had assumed that the echidna egg tooth had this same layer (Green, 1930; Hill & De Beer, 1950). However, recently it was discovered that both the platypus and echidna have lost a number of genes involved in tooth formation and echidnas have lost additional enamel producing genes compared to the platypus (Zhou et al., 2021). This is unsurprising since whilst both platypus and echidnas possess an egg tooth, once the echidna egg tooth falls out they are edentulous whereas platypuses develop cheek teeth in the nestling stage and these do contain enamel (Davit-Béal et al., 2009). The short-lived nature of the echidna egg tooth and the soft, leathery form of the egg means that a hard enamel layer is unlikely to be necessary for hatching. The presence of enamel may, therefore, depend on the chemical composition of the eggshell, or vice versa.

In further contrast to other mammals, the trabecular bone of the echidna and platypus premaxilla was fused to the egg tooth (ankylosed) rather than being held in a socket by a periodontal ligament as occurs in all other mammals examined to date. The rapid projection of the tooth between days 4 and 6 and the absence of any invagination of the epithelium, means that a root was not evident as roots form from invaginations of the epithelium (Thesleff & Juuri, 2012). In the absence of invagination in the echidna, the dentine made direct contact with the bone for attachment. A fused tooth is sturdier than a tooth attached via soft tissue (Jenkins & Shaw, 2020). An ankylosed connection can also likely be established faster than a thecodont attachment. This would be an advantage for the echidna since they have only a brief period of time to develop a structure strong enough to allow them to hatch from their egg. Acrodont attachments are also evident in squamate egg teeth, with the dentine attached to the bone via an attachment tissue (Hermyt et al., 2020). Before loss of the egg tooth, large numbers of TRAP positive clast cells were observed within the egg tooth, lining the dentine at the base of the tooth. These cells appeared to be odontoclasts, as they were observed adjacent to regions of thinned dentine, and thereby could break the connection between the tooth and premaxilla. Similar multinucleated cells were observed in the platypus, highlighting a conserved mode of tooth loss in monotremes. Removal of the dental pulp was then completed by programmed cell death.

The premaxilla extended up and appeared to fuse with the os caruncle, thereby connecting the egg tooth and caruncle into one functional unit. The premaxilla was one of the first bones in the head to ossify, confirming its important support role for both the egg tooth and caruncle. In addition to the formation of the os caruncle in the mesenchyme, the epithelium overlying this structure was cornified, with a more complex stratified epithelium, with similarities to that observed in birds and turtles (Alibardi, 2020). This suggests that while the mesenchymal os caruncle may be a novelty in monotremes, the epithelial caruncle may be homologous to that of reptiles, suggesting that the common amniote ancestor of reptiles and mammals had both of these characteristics. This would suggest that reptiles without caruncles have lost these as they came to rely more heavily on the egg tooth itself. Alternatively, the thickened epithelial caruncles might have evolved independently in the different lineages (turtles, crocodilians, birds, sphenodon, monotremes). Loss or gain of the caruncle and egg tooth is likely to be closely linked with changes in shell composition.

Overall, the results from this study support previous suggestions that the monotreme egg tooth development pathway is conserved through an ancestral type of tooth development. The echidna egg tooth was very similar to the primary teeth of some reptiles. Both are ankylosed teeth that develop rapidly through evagination of the oral epithelium rather than invagination, and both have little to no enamel. Together, these results highlight the similarities between reptiles and monotremes and show conserved traits from a common amniote ancestor over 315 million years ago.

## Materials and Methods

### Animals

Fetal and pouch young short-beaked echidna (*Tachyglossus aculeatus*) samples were collected from a research breeding colony at Currumbin Wildlife Sanctuary. Fetal samples were dated based on the date of egg laying (oviposition), with samples varying in age from day 4 to day 9 after oviposition (where hatching occurs on day 10) (day 4 n=1, day 6 n=1, day 7.5 n=1, day 8 n=3, day 9 n=2). Pouch young samples were dated based on the day of hatching (day 0 py) (day 0 n=2, day 2 n=2, day3 n=1, day 4, n=1 day 11, n=1, d50 n=1). All samples were fixed in 4% paraformaldehyde overnight, washed twice in phosphate buffered saline and stored in 70% ethanol at 4°C. All samples were approved by the University of Queensland Animal Ethics Committees and followed the Australian National Health and Medical Research Council guidelines (2013). Historic slides of a platypus at day 2 hatching (M44 (WW) 16.75mmm collected in 1899) were photographed from the Hill Collection, Museum fur Naturkunde, Leibniz Institute for Research on Evolution and Biodiversity, Berlin.

### Micro-CT Scanning and 3-D reconstructions

One of the fetuses had already been collected and processed for histology before the commencement of this project so was unable to be micro-CT (micro computed tomography) scanned. All remaining echidna fetuses and pouch young samples were scanned by micro-CT. Micro-CT scanning of the echidna fetuses and the d11 py was performed with a phoenix nanotom m (Waygate Technologies, Huerth, Germany) operated using xs control and phoenix datos|x acquisition software (Waygate Technologies). An X-ray energy of 35-40 kV and 300 μA was used. Scans were conducted using a molybdenum target to maximize contrast from the soft tissue specimens. The voxel resolution was optimised to the size of the specimen and varied from 3.1 to 11.1 μm. The number of X-ray projections collected through a full 360-degree rotation was also optimized to the size of the specimen, varying from 599 to 1798 projections leading to scan times of 5 to 15 min in a fast scan mode (Table 1). The data was exported as 16-bit volume files for imaging and analysis. All remaining pouch young samples were scanned using a Scanco micro-CT scanner and images analysed using Amira (ThermoFisher Scientific, Waltham, MA, USA).

**Table 1:** Micro-CT scan data for 7 echidna fetuses.

| Sample description | Scan time (min) | Target | Voltage (kV) | Current (μA) | Resolution (μm) | Projections |
| --- | --- | --- | --- | --- | --- | --- |
| d4 unstained | 5 | Molybdenum | 35 | 300 | 3.1 | 599 |
| d4 stained | 12 | Molybdenum | 40 | 300 | 3.111 | 1438 |
| d6 unstained | 12 | Molybdenum | 35 | 300 | 4 | 1438 |
| d6 stained | 12 | Molybdenum | 40 | 300 | 3.502 | 1438 |
| d7.5 stained* | 12 | Molybdenum | 35 | 300 | 4 | 1438 |
| d8 stained* | 12 | Molybdenum | 35 | 300 | 4 | 1438 |
| d8 stained* | 13 | Molybdenum | 35 | 300 | 3.1 | 1558 |
| d8 unstained | 13 | Molybdenum | 35 | 300 | 4.001 | 1558 |
| d8 stained | 2x10 | Molybdenum | 40 | 300 | 4 | 1199 |
| d9 stained* | 15 | Molybdenum | 35 | 300 | 4 | 1798 |

\*previously stained and washed, stored in 70% ethanol

Three fetuses were scanned twice, first unstained to detect mineralised tissues and secondly after being stained with 1% iodine in 100% ethanol overnight to increase the contrast of non-mineralised tissue. All other fetal samples were scanned once after being previously stained with 1% iodine in 100% ethanol overnight and then cleared with sodium thiosulfate before being stored in 100% ethanol. After the fetuses were iodine and micro-CT scanned, they were washed in 70% ethanol to remove any excess iodine stain and then incubated in sodium thiosulfate to clear the iodine colouration before paraffin embedding and sectioning. Pouch young samples were scanned twice.

3D image analysis and segmentation were performed in Avizo v9.7.0 (ThermoFisher Scientific) via the MASSIVE platform (Goscinski et al., 2014).

### Histology

After micro-CT scanning, fetuses were sectioned at a thickness of 5 μm through the sagittal plane on a Rotary CUT 4060 microtome (Microtec Laborgerate, GmBH, Walldorf Germany) and dried overnight in a 40°C oven. Every third slide was then de-waxed and rehydrated through a graded ethanol series to distilled water (dH_2_O) and stained with haematoxylin and eosin (H&E) using standard techniques. From the H&E-stained slides, the tooth region was identified, and in this region, every second remaining slide was stained with a Picro-Sirius Red and Alcian blue trichrome stain for histological examination of bone, cartilage and dentine using standard techniques. The remaining slides were kept for immunofluorescence

### TRAP staining

In order to visualise odontoclasts, dewaxed and rehydrated sections were tartrate-resistant acid phosphatase (TRAP) stained as previously described (Anthwal et al., 2017). Briefly, rehydrated slides were incubated in filtered pH 5.32 acetate buffer containing 1 mg/ml Naphthol-AS-TR-phosphate (Sigma), 100 mM Sodium Tartrate (Sigma-Aldrich, Merck, Darmstadt Germany), and 1 mg/ml Fast Red TR Salt (Sigma-Aldrich) at 37°C until colour reaction was complete. Slides were then counterstained with Mayer’s Haematoxylin (Sigma) for 10 seconds, and blued in tap water, and mounted in aqueous mounting medium (Aquatex, Sigma-Aldrich)

### Immunofluorescence

Sections used for immunofluorescence were first de-waxed in histolene and rehydrated as above, with a final wash in dH_2_O. To detect protein expression of sonic hedgehog (SHH), collagen type II (Col2), cytokeratin 14 (K14), cytokeratin 8 (K8), and Loricrin, sections were first placed in the antigen retrieval buffer 0.01 M Sodium citrate pH10 (w/v) and incubated in a preheated water bath at 95°C for 20 min before being left to cool at room temperature for 40 min. After cooling, the slides were washed in 1X phosphate buffered saline, (1X PBS) containing 1.37 M NaCl, 27 mM KCl, 18 mM KH_2_PO_4_ and 100 mM Na_2_HPO_4_. Following the 1x PBS wash, sections were incubated in 0.3% Sudan Black (w/v, ProSciTech, Qld, Australia) in 70% ethanol for 10 min before being immersed briefly in fresh 70% ethanol and then washed for 5 min in 1X PBS. Sections were then blocked with 10% (v/v) normal goat serum in 1x PBS for 1 h at room temperature. After blocking, the sections were incubated overnight (16 h) in a humid chamber at 4°C with either rabbit SHH antibody (2 μg/ml; ab19897; Abcam, Cambridge, UK), mouse Col2 antibody (1/50 dilution; II-II6B3; Developmental Studies Hybridoma Bank, Iowa City, IA, USA), mouse K8 antibody (1/50 dilution; TROMA-1; DSHB), or co-labelled with mouse anti K14 (0.625μg/ml; ab7800; Abcam) and rabbit Loricrin antibody (1μg/ml; 905101; BioLegend, San Diego, CA, USA). Sections were then washed three times in PBS containing 0.5% Tween 20 (v/v) then incubated for 45 min with goat anti-rabbit Alexa Fluor Plus 555 secondary antibody (A32732, Invitrogen, ThermoFisher Scientific) at either 1:300 dilution (SHH), goat anti-rabbit Alexa Fluor 568 (A11011, Invitrogen) at 1:400 dilution (Loricrin), or goat anti-mouse Alexa Fluor 488 (A32723, Invitrogen) at 1:400 (COL2, K8, K14). Nuclei were counterstained with 4′,6-diamidino-2-phenylindole (DAPI; D9542; Sigma-Aldrich) and the slides mounted with media (homemade, or Fluroshield, Abcam) and kept in the dark until visualisation. Negative control sections were submitted to the same procedures except that the first antibody was replaced by the relative isotype control at the same concentration as the primary antibody. The specificity of all antibodies was confirmed by staining in positive and negative control tissues and lack of staining in the isotype controls. Images were acquired on a Nikon (Minato City, Japan) A1R spectral confocal microscope housed at the University of Melbourne, or a Zeiss Apotome fluorescence microscope housed at King’s College London. Visualisation settings were initially optimised to eliminate background using the isotype control on each slide. These settings were then used to visualise the remaining positive sections on the slide.

## Statement of Ethics

The University of Queensland Animal Experimentation Ethics Committee approved all sampling for echidnas, in accordance with the National Health and Medical Research Council of Australia Guidelines (*Australian code for the care and use of animals for scientific purposes*, 2013; 2013).

## Author contribution statement

JCF, NA, AST, ARE and MBR designed research; JCF, AB and NA performed the research; JCF, SDJ, MBR and MP collected tissues, JCF, AB, NA, ARE, AST and MBR analysed data; and JCF, AB, NA, AST and MBR wrote the paper. All authors read and approved the final version of the manuscript.

## Acknowledgements

Particular thanks to the veterinary staff and keepers at Currumbin Wildlife Sanctuary for echidna sample collection. Thanks to the Melbourne TrACEES (Trace Analysis for Chemical, Earth and Environmental Sciences) platform for access to the micro-CT scanner and Dr. Jay Black for technical support. This work was supported by the MASSIVE HPC facility (www.massive.org.au). Thanks also to Dr Peter Giere for access to the JP Hill collection housed at the Museum fűr Naturkunde, Berlin.

---

**Figure 3 - figure supplement 3:**
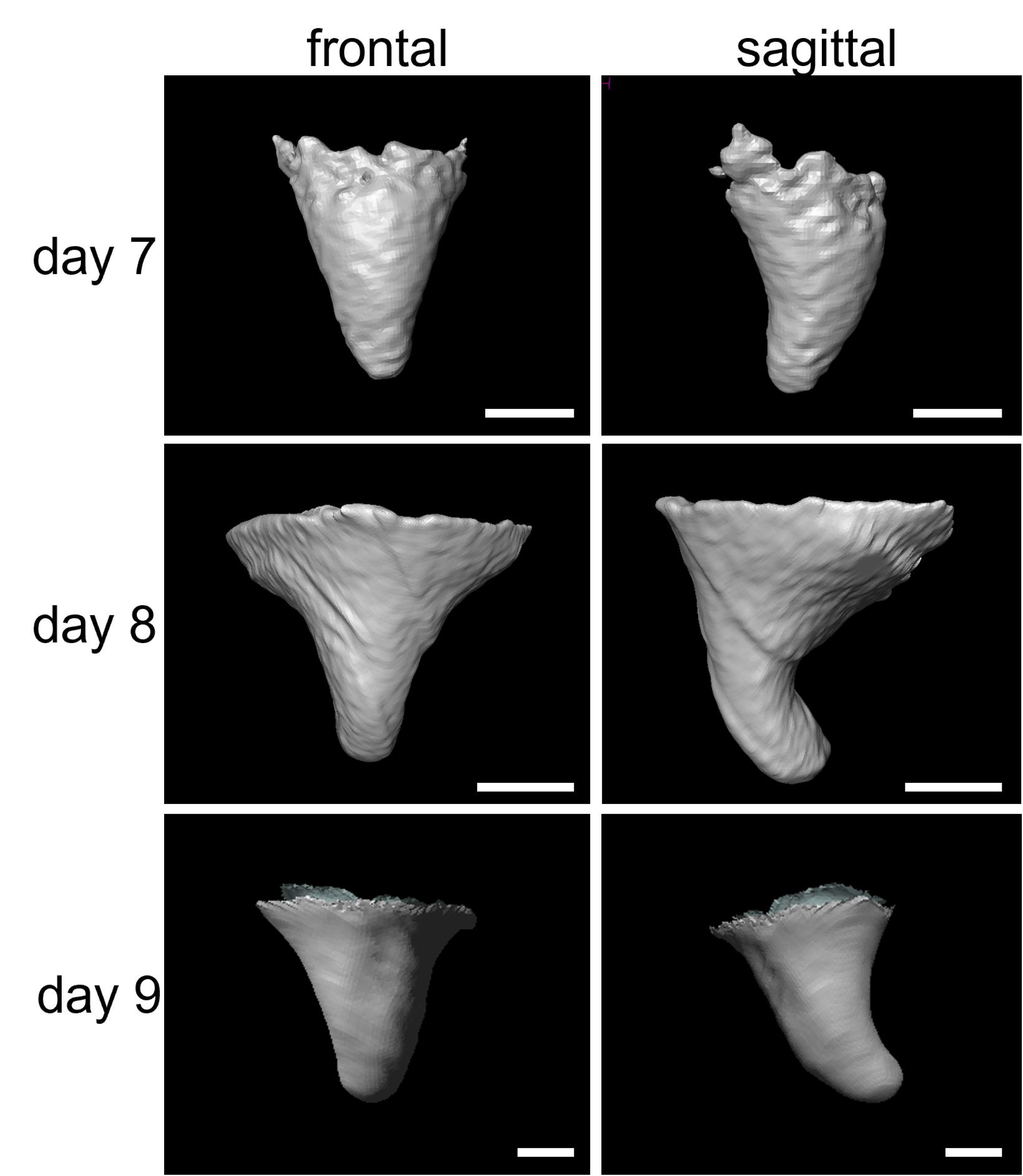
3-dimensional models of the echidna egg tooth through the frontal and sagittal planes at different days of development. Micro-CT volume rendering of the mineralised tissue. Scale bars= 100 μm.

**Figure 5- figure supplement 5:**
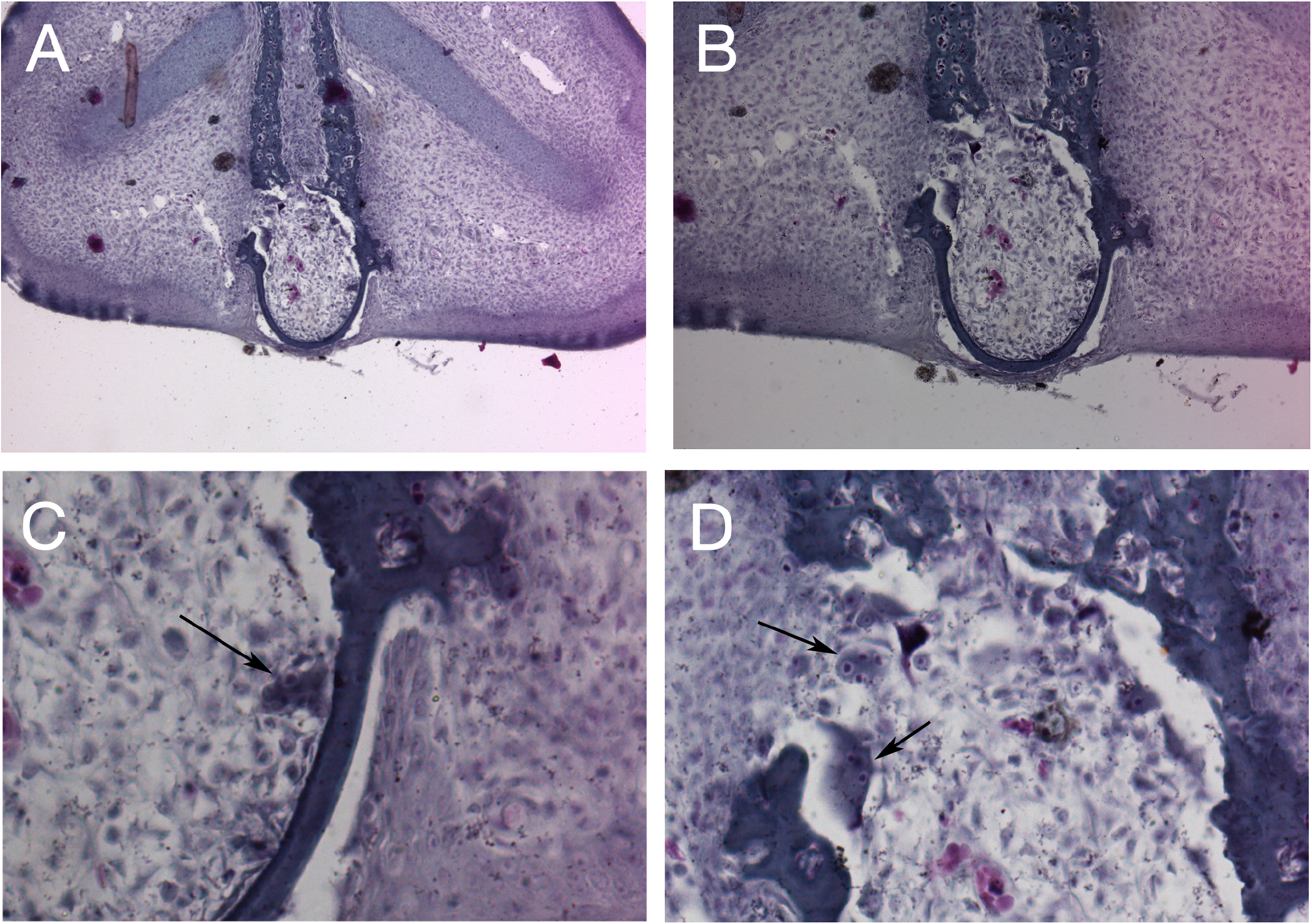
Platypus (*Ornithorhynchus anatinus*) 2 days post hatching. A-D: Frontal views of egg tooth. A,B: The egg tooth has started to detach from the premaxilla with a break evident on the LHS. C,D: Multinucleated cells are evident inside the dental pulp lining the dentine (C), and at the border between the tooth and premaxilla in the regions of bone loss (D).

